# THE INFLUENCE OF FOOD ABUNDANCE CHANGE ON SEARCH AND ATTACK BEHAVIOR OF INSECTIVOROUS BIRD IN THE MONTE DESERT, ARGENTINA

**DOI:** 10.1101/2025.02.20.639347

**Authors:** Carolina Guerra-Navarro, Victor R. Cueto

## Abstract

Bird foraging is a combination of two behaviors, the search movements to locate preys and the attack maneuvers to catch them. However, the first behavior is poorly studies. Hear we determine the searching movements and attack maneuvers of three species of insectivorous birds in the central Monte desert, Argentina and evaluate if these two components of the foraging behavior change as a function of interannual variation in the availability of food resources. During three years we studied in the Ñacuñán Reserve (Mendoza, Argentina) the search movements and attack maneuvers of three insectivorous bird species in the foliage of *Neltuma flexuosa* trees and simultaneously evaluated the abundance and biomass of their prey. We found that arthropod abundance increased in the last year, although biomass was low, due to the greater number of small arthropods (less than 0.2 mg) in that year. During the first and second year the main capture maneuver of Grey-crowned Tyrannulet (*Serpophaga griseicapilla*) was sally-hover and it moved among the foliage using flight movements, but in the third year a greater use of gleaning maneuvers was observed and its main foraging movements were jumping on the branches.

Greater Wagtail-tyrant (*Stigmatura budytoides*) used both sally-hovering and gleaning maneuvers in the first and second years, however it used glean more frequently and performed more jumping movements in the third year Ringed Warbling-Finch (*Microspingus torquata*) showed very stereotyped behavior throughout the study period, using only gleaning and jumping as foraging strategies. The increase of small prey in the last year could have determined the changes in attack maneuvers and searching movements of Grey-crowned Tyrannulet and Greater Wagtail-tyrant. The gleaning maneuver might be more suitable for capturing small prey, probably because birds detect them better at close range, and moving with short movements such as jumping might allow them to explore branches and foliage in greater detail. For Ringed Warbling-Finch, given the characteristics of its feeding behavior, the above-mentioned changes did not influence the way it search and attack prey. The interannual changes observed in searching and capture behavior provide valuable information about ecological flexibility of insectivorous birds that may be advantageous when they confront variable climates and food fluctuations such as in desert environments.

## Introduction

The first component of foraging behavior is the prey search (Remsen and Robinson 1990). This behavior includes the movements involving the position changes that birds use to locate their food (Robinson and Holmes 1982). Thus, “foraging” begins with all movements made prior to locating the food or substrates that conceal it and ends once it is located and the prey attack has begun (Remsen and Robinson 1990). Search movements exclude those that the bird performs to catch its prey, which are part of the attack maneuvers. These movements are represented by “foraging speed” (i.e., the number of total movements of the bird per unit time), and reflects the intensity of foraging effort by birds (Robinson and Holmes 1982, Dobbs et al. 2007). However, search behavior has received little attention in the literature (Lyons 2005). The movements that a bird performs when are searching can be classified into different categories, for example: jumping, running, climbing (using the tail or not), gliding, flapping, and flying (Remsen and Robinson 1990). Studies of searching behavior commonly analyze the rates of each of these movements between perches (branches or substrates that the bird uses for grasping) and the time interval between movements (Robinson and Holmes 1982, Holmes and Recher 1986, Lyons 2005, Chen et al. 2011, Oyugi et al. 2012). These variables reflect to what extent a given area is explored, in what manner and how much a bird moves from perch to perch in its search for prey (Robinson and Holmes 1982). Considering the different types of movements some studies suggest that jumps reflect movements within a feeding patch and flights reflect movements between patches (Oyugi et al. 2012). In addition, different types of jumps could imply different areas explored, for example jumps on the same perch (i.e., on the same branch where the bird was located) implies a smaller area explored than a jump in which the bird changes perch.

The second sequential component of bird foraging behavior corresponds to the attack on prey (Remsen and Robinson 1990). The attack is considered as the sequence that goes from the moment the food item is visualized by the bird until the capture attempt is made by means of a particular maneuver or movement (Remsen and Robinson 1990). Attack maneuvers are mainly determined by the bird’s morphology, but factors extrinsic to the bird may also play a role, such as changes in the availability of different types of prey (Martin and Karr 1990, Böhm and Kalko 2009). The foraging behavior of birds may undergo interannual variation in response to changes in the abundance of food resources and/or variations in environmental conditions (Szaro et al. 1990, Bell and Ford 1990). Interannual change have been observed in behavioral variables such as foraging site - plant species and substrates - (Wagner 1981, Hejl et al. 1990, Miles 1990, Szaro et al. 1990, Unno 2002, Cueto et al. 2016) and in the use of different attack maneuvers (Ford et al. 1990, Szaro et al. 1990). This ability of birds to respond to changes in the food resource or to the presence of new resources is determined by the ecological flexibility of individuals (Greenberg 1990).

Our aim is to determine the searching movements and attack maneuvers of three species of insectivorous birds in the central Monte desert, Argentina. Also, we evaluate if these two components of the foraging behavior change as a function of interannual variation in the availability of food resources.

## Methods

### Study area

The study was carried out in the Biosphere Reserve of Ñacuñán (34º 03’ S, 67º 54’ 30’’ W), located in the central Monte desert, Mendoza, Argentina. The landscape of the reserve is mainly a mesquite open woodland intersected by variably sized tracts of creosotebush shrubland. The mesquite woodland is a matrix of non-thorny tall shrubs (mainly *Larrea divaricata*), with thorny trees (*Neltuma flexuosa* and *Geoffroea decorticans*) and tall shrubs (*Capparis atamisquea* and *Condalia microphylla*). Nonthorny tall shrubs (*Larrea cuneifolia*) dominate the creosotebush shrubland, with scarce cover of thorny trees and tall shrubs (Marone 1991). The open woodland has higher vertical complexity (three vegetation strata) than the shrubland (two strata) (Marone 1991). The climate is dry and highly seasonal. Mean annual rainfall is 342 mm, and more than 75% of the annual rainfall occurs in the warmer months (October–March). Therefore, the summer and spring are hot and rainy and the winter and autumn are cold and dry.

### Search and attack behavior sampling

We recorded the searching and prey attack behavior of Greater Wagtail-tyrant (*Stigmatura budytoides;* 11 g) and Grey-crowned Tyrannulet (*Serpophaga griseicapilla;* 5.5 g) belonging to the family Tyrannidae, and Ringed Warbling-Finch (*Microspingus torquata;* 10 g) belonging to the family Thraupidae (this latter species shifts from the foliage-foraging guild to the graminivore guild during the non-breeding season, Lopez de Casenave et al. 2008). Greater Wagtail-tyrant and Ringed Warbling-Finch are resident birds in the study area (Marone 1992); Grey-crowned Tyrannulet is a short migrant bird, present at the study area during spring and summer (Cueto et. al 2008). We gathered foraging data during three breeding-seasons (October-March): 2006-2007, 2007-2008 and 2008-2009. During each sampling period, we systematically walked through the three plots (10 hectares each) from sunrise to midday and from the afternoon until sunset, except on rainy or too windy days. We avoid walked through the same area twice on the same day to avoid repeating the observations of the same individual birds.

Each time a bird was observed searching for food, its behavior was recorded until it was lost from sight. Data were recorded for individuals that were engaged in foraging activities, disregarding those that were resting, singing, or engaged in any other activity that might affect feeding rates (Robinson and Holmes 1982). If an individual was engaged in any of these other activities within a feeding sequence, the start and end of that activity (e.g., singing in the middle of the feeding sequence) was recorded and then subtracted from the final sequence. Observations were recorded on a digital recorder and the following variables were considered: bird species, prey attack maneuver, and searching movements within the foliage of an individual of *Neltuma flexuosa* tree. We restricted the observation only in this tree species because the three bird species select *Neltuma flexuosa* as foraging site (Guerra-Navarro and Cueto 2024) and to avoid the effect of vegetation architecture of other plant species in the foraging behavior of the insectivorous species. Search movements were categorized (based on Remsen and Robinson 1990) into: (1) jumping: movement within the same branch (perch) by jumping without using its wings, (2) change of perch by jumping: movement from one branch to another by jumping, without using wings, (3) change of perch by flying: movement from one branch to another by flying. Recorded data were transcribed into spreadsheets using a stopwatch, and the time that each sequence of observed behavior lasted was determined. The variables related to searching behavior were defined as the number of a given type of movement per minute (Robinson and Holmes 1982). Thus, the new variables defined were: jumping rate, perch change per jump rate, perch change per flight rate and search speed (total movement rate). These rates were calculated for each observed sequence of each individual. If the same individual was observed during the same sampling day or if it changed of tree, the movement rates were averaged to obtain a single data corresponding to that sampling day. The bird species studied are small and move very fast through the foliage (so it is difficult to obtain feeding sequences of more than 30 seconds) and it is rarely possible to follow the individual from the time it climbs up to the time it leaves a tree. Due to these drawbacks, the recorded sequences were classified into (a) complete sequences: feeding sequences that began with the arrival of a bird at a tree to feed until the individual left it, and b) incomplete sequences: feeding sequences where the arrival and/or departure of the bird could not be observed. Complete foraging sequences and some of the incomplete sequences were considered for the estimation of movement rates and statistical analysis. For an incomplete sequence to be incorporated in the analysis, it was decided to use a “minimum time” criterion to include in the data base. This “minimum time” was estimated for each bird species as the average of the time duration of all complete sequences. This criterion proved to be conservative compared to other studies where foraging sequences with minimum times of 10, 20 or 30 seconds were considered in the analysis (e.g., Chen et al. 2011, Robinson and Holmes 1982, Holmes and Recher 1986). Further, choosing a different minimum time for each species as a criterion for defining which sequences to use, contemplates differences in the foraging behavior of the species (Salewski et al. 2003).

Prey-attacking maneuvers were sorted into three categories (based on Remsen and Robinson 1990): (1) glean: when a perched bird captures food from the surface of a nearby substrate; (2) sally-hover: when a bird captures food from the surface of a substrate while in flight; (3) sally-strike: when a bird flies from a substrate to capture a prey in the air. To determine the frequency of use of attack maneuvers, the methodology of Airola and Barrett (1985) was used, because the data to be analyzed are sequential observations (for more details see Guerra-Navarro and Cueto 2024 and reference there in).

Data obtained in the three plots were pooled, since no differences were observed in the searching and prey-attacking maneuvers of the birds.

### Arthropods sampling

We used the “branch clipping” method (Cooper and Whitmore 1990, Johnson 2000) to evaluate the arthropod abundance and biomass in the *Prospis flexuosa* foliage. The arthropods sampling was carried out in the same period of the observations of birds foraging behavior. The sampling was carried out between 10:30 am to 1:30 pm. We randomly selected 10 individuals of *Neltuma flexuosa* in every plot. We collected an arthropod sample enclosing one branch (approximately 50 cm) into a plastic bag as quickly as possible. Then we clipped the branch with shears and fumigated the sample with insecticide. For each sampled branch we recorded its length, the longest branch diameter and the perpendicular diameter to the longest. We quantified the number of arthropods recovered from each sampled branch up, these arthropods were identified to Order or Family level and their body length was measured with a micrometer under a binocular microscope; we only considered the arthropods larger than 1 mm following others studies (Wiens et al. 1991, Diaz et al. 1998). The dry weight (W, mg: 60°C, 48 h) of arthropod was estimated from the body length (L, mm) with equations derived from weights and lengths measured on a subset of the samples collected at the study site (following Hódar 1996, see Guerra-Navarro and Cueto 2024). We used different equations based in the order/family level and we take into account morphological characteristics as shape and body toughness (Hódar 1996). Equations can be found in Guerra-Navarro and Cueto (2024). For each sample we estimated the volume for the clipped branch assuming as cylinder-shaped. We then estimated the abundance and biomass of arthropods per m^3^. Arthropod abundance and biomass data from three plots were pooled as no differences were found between them.

### Statistical analysis

Search speed was calculated as the sum of the three types of searching movements (branch jump, perch changes per jump, and perch changes per flight) per minute for each individual. Inter-annual variation in foraging speed was analyzed using a two-factor ANOVA (factors: bird species and year). Data were transformed using the square root transformation of the data (x’ = √ (x + 0.5)) to meet the assumption of homogeneity of variance among variables (Zar 1996).

Inter-annual variations in the rates of the different searching movements of the birds were analyzed by means of a multivariate analysis of variance (MANOVA). In this analysis, the rates of searching movements were the variables measured on each individual bird and the treatment corresponds to the years. The movement rates included in this analysis were: jumping rate, perch change per jump rate and perch change per flight rate. The search speed variable was analyzed separately because it is a composite variable of those used for the MANOVA. When the MANOVA was significant, the recommendation to perform a Discriminant Analysis (Johnson and Field 1993) was followed to evaluate the effect of the individual variables, i.e. which variable contributed to the differences between sampling years. The square root transformation of the data (x’ = √ (x + 0.5)) was used to meet the assumption of homogeneity of variance for each variable (Zar 1996). Barlett’s test was used to evaluate the hypothesis of homogeneity of covariance matrices (Morrison 1976).

For each bird species, a test of independence was performed to evaluate whether the frequency of use of the different attack maneuvers is independent of the year. For each test of independence, the requirement that the average of the expected frequencies be greater than 10 in order to work with a type I error equal to 0.01 was checked (Zar 1996). To evaluate the degree of dependence of the maneuvers on the years, Pearson’s Chi-square statistic was used. In addition, to analyze the nature of the associations between the use of maneuvers and years, Pearson’s standardized residuals were calculated, which provide information on the strength of the association (Agresti 2002). The criterion that if a standardized Pearson residual exceeds an absolute value of 2 indicates a lack of fit to the null hypothesis of independence in that cell was followed (Agresti 2002).

To analyze the effect of sampling year on arthropod abundance (number of individuals per branch), a generalized linear model using the Negative Binomial probability distribution was used. This distribution was adequate because the data set presented a strong “overdispersion” (Quinn and Keough 2002). This “overdispersion” usually appears when the data come from counts (in this case: number of arthropods per unit volume) and present a variance greater than the mean (Agresti 2002), therefore it is incorrect to use a Poisson distribution (which assumes a mean equal to the variance). Because the volume per branch varied from one sample to another, the volume of each branch was used as an offset variable, becoming a constant that is added to the model (Littel et al. 1996, Zuur et al. 2009); in this case the offset constant is the natural logarithm of the volume of each branch. This offset variable can only be used for Poisson, Negative Binomial and Geometric distributions (Zuur et al. 2009). Thus, the variable to be evaluated was the abundance of arthropods per sampled branch; this is more realistic since it allows assigning to the model the volume value corresponding to each unit (Mangeau and Videla 2005), in this case each branch. To assess the significance of the “year” factor on arthropod abundance, a log likelihood ratio test (log likelihood ratio test, Zuur et al. 2009) was performed comparing the fitted model with a null generalized linear model (without the “sampling year” factor incorporated).

To analyze the effect of sampling year on arthropod biomass, analyses were performed considering the variable biomass per volume (mg/m3), but due to the variability of the data and the lack of compliance with the assumption of homogeneity of variance, a generalized least squares (GLS) analysis was performed. The GLS analysis is an extension of the generalized linear models, which consists of a weighted linear regression that allows modeling the heteroscedasticity of the data generated by the factors that are contributing to the variance inequality (Pinheiro and Bates 2000, Zuur et al. 2009); in this case due to the sampling year factor. The modeling of the residual variance at the different levels of the “year” factor was performed using the varIdent function (Zuur et al. 2009). To compare the model without the incorporated variance structure with the model with modeled heteroscedasticity, a log likelihood ratio test (log likelihood ratio test, Zuur et al. 2009) was performed. In addition, the model was graphically validated using the standardized (normalized) residuals following the guidelines of Zuur et al. (2009). To evaluate the significance of the “year” factor on arthropod biomass, a log likelihood ratio test was performed comparing the fitted model with a null generalized linear model (without the “sampling year” factor).

In addition, based on the individual mass of the arthropods collected, mass ranges were established and the frequency of individuals in each of these ranges was determined for the three years studied. An Independence Test was used to evaluate whether the frequencies of the different mass ranges were independent of the years. The requirement that the average of the expected frequencies be greater than 10 in order to work with a Type I error equal to 0.01 (Zar 1996) was corroborated. The Pearson Chi-square statistic was used to evaluate the significance of the independence of the mass categories with the years.

All statistical analyses were performed with Infostat (2010), except for the linear models for arthropod abundance and biomass, for which the AED, MASS and NLME packages of the R software (R Core Team 2012) were used.

## Results

### Search speed and searching movements

The average times of complete sequences for each bird species were: Grey-crowned Tyrannulet: 25.7 s, Greater Wagtail-tyrant: 47.4 s, and Ringed Warbling-Finch: 54.5 s. After applying the selection criterion to incomplete sequences, the total number of independent sequences for the estimation of movement rates for the three bird species was 380, totaling 543.8 minutes of feeding record.

For the search speed no significant interaction was found for the factors bird species and year (F_4, 371_ =1.26, P =0.28). The three bird species had different foraging speed (F_2, 371_ = 25.7, P<0.0001; Fig. 1). Individuals of Grey-crowned Tyrannulet showed the lowest foraging speed, whereas individuals of Greater Wagtail-tyrant performed the most searching movements per minute. For each species, search speed was not different among the three years studied (F_2, 371_ =1.39, P =0.25; Fig. 1).

**Figure 1.**
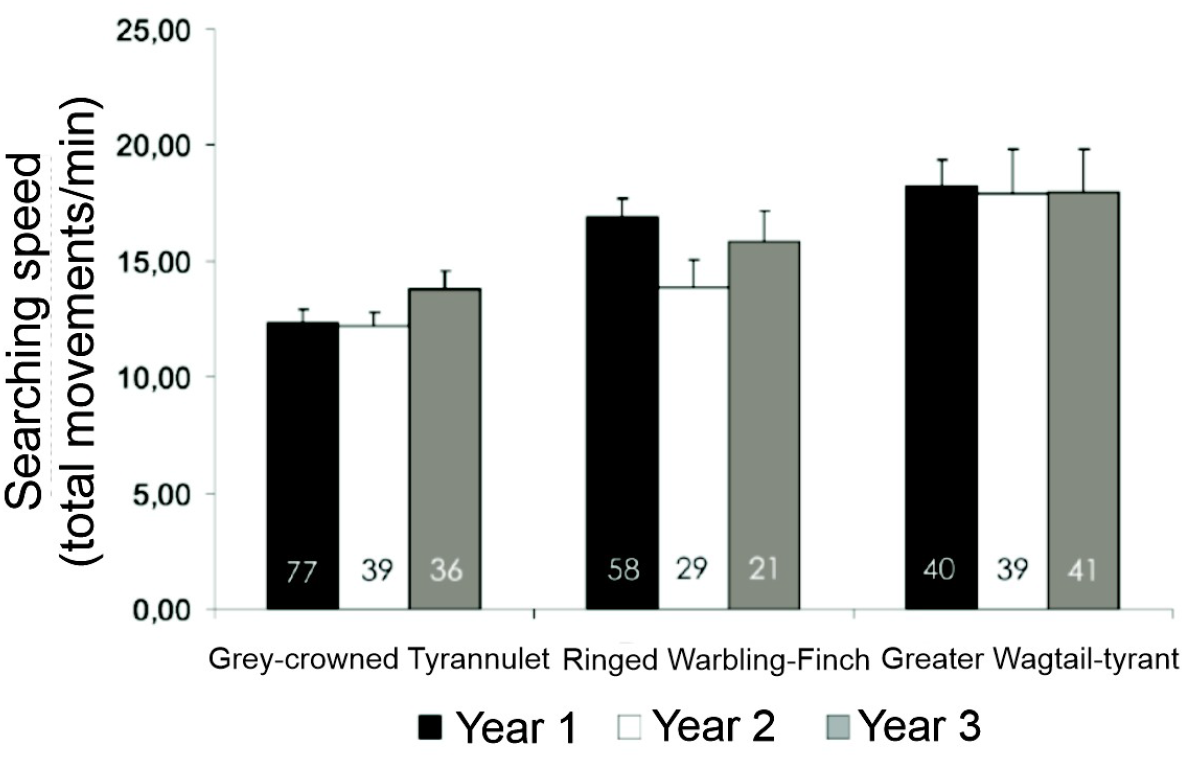
Searching speed (+ EE) for the three insectivorous bird species during three years in the central Monte desert, Argentina. Movements correspond to the sum of perch jumps, perch changes per jump and perch changes per flight for each individual. Species factor: F_2, 371_ = 25.7, P<0.0001; year factor: F_2, 371_ =1.39, P =0.25; species-year interaction: F_4, 371_ =1.26, P =0.28. N is indicated on each bar of the graph.

MANOVA for searching movements indicate that there were significant differences among the years for the three species (Fig. 2). Grey-crowned Tyrannulet searching movement rates was significantly different for the first two years compared to the third (Fig. 2). When performing a Discriminant Analysis more than 99% of the variation was explained by the first discriminant function (Table 1); that was characterized by perch changes per flight for the first two years and by perch changes per jump during the third year (Table 1 and Fig. 2). Greater Wagtail-tyrant searching movement rates in the first year was significantly different from the multivariate response in other years (Fig. 2). Discriminant Analysis indicate that the first discriminant function (which explains more than 95 % of the variation) was characterized by perch changes per flight among years (Table 1). The greatest number of changes of perch per flight was in the first year (Fig. 2). In the last two years, this variable became the variable least used by Greater Wagtail-tyrant for searching preys (Fig. 2). For Ringed Warbling-Finch significant differences were found between the first and the third years (Fig. 2). The Discriminant Analysis showed that for the first discriminant function (which explains 82 % of the variation) the variable that contributes most to the differences between these two years was the rate of perch change per flight (Table 1). The second discriminant function (which explains 14 % of the variation) indicate that during the last year, Ringed Warbling-Finch also increase the rate of branch jumping (Table 1).

**Table 1.** Coefficients of the search movement variables for each discriminant function for three insectivorous bird species of the central Monte desert, Argentina. Coefficients were not standardized since the variables have the same units. The largest coefficients for each discriminant function are indicated in bold. For description of search movement variables see Methods.

|  | ***Discriminant function*** |  |
| --- | --- | --- |
|  | **1** | **2** |
| **Grey-crowned Tyrannulet** |  |  |
| Constant | -0.28 | 0.52 |
| Jump rate | -0.39 | **1.64** |
| Perch change per jump rate | **1.19** | -0.5 |
| Perch change per flight rate | -0.71 | -0.52 |
| **Explained variance (%)** | **99.5** | **0.5** |
| **Ringed Warbling-Finch** |  |  |
| Constant | -3.16 | -3.9 |
| Jump rate | -0.24 | **1.02** |
| Perch change per jump rate | 0.1 | 0.5 |
| Perch change per flight rate | **1.99** | 0.11 |
| **Explained variance (%)** | **82.6** | **14.4** |
| **Greater Wagtail-tyrant** |  |  |
| Constant | -1.44 | -1.99 |
| Jump rate | -0.27 | **1.51** |
| Perch change per jump rate | -0.21 | -0.61 |
| Perch change per flight rate | **1.37** | 0.51 |
| **Explained variance (%)** | **95.3** | **4.7** |

**Figure 2.**
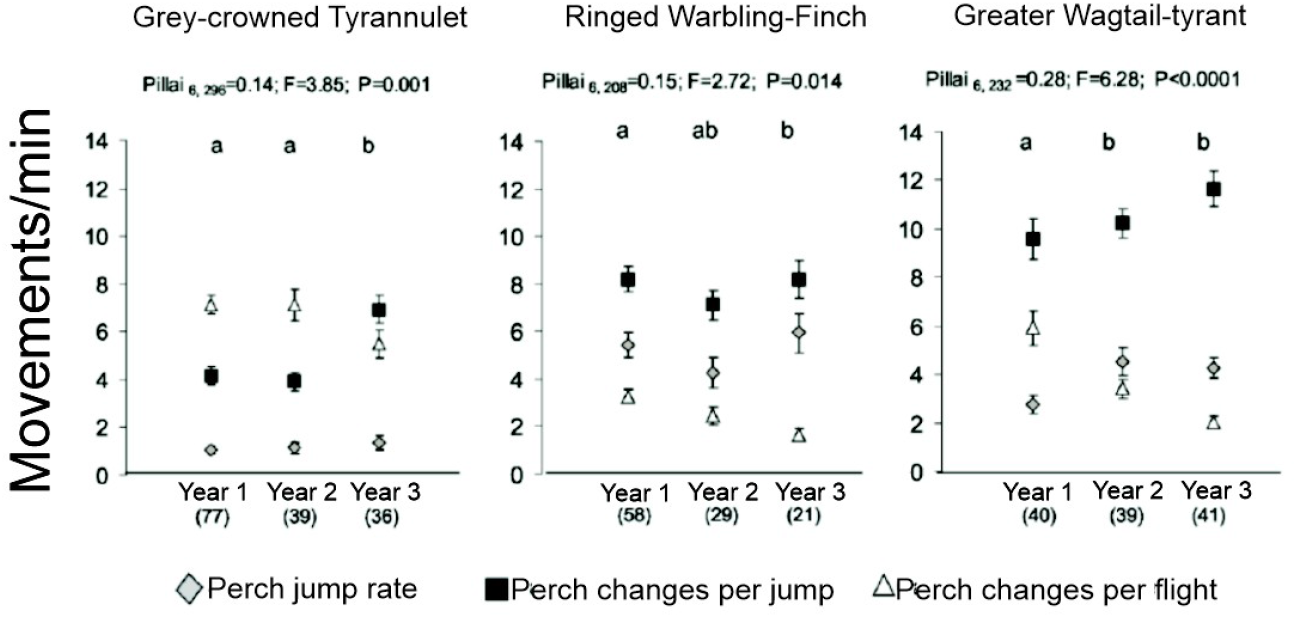
Search movement rates (number of movements per minute + EE) for three insectivorous bird species during three years in the central Monte desert, Argentina The value of the Pillai trace statistic, the F statistic and the P value for the MANOVA are indicated. Comparisons between years were performed with the Bonferroni-corrected Hotelling Test (different letters indicate significant differences, P < 0.05). The number of independent sequences for each period is indicated in parentheses.

Prey-attack maneuvers was year-dependent only for Grey-crowned Tyrannulet (X^2^_4_ = 9.57, P=0.04, Fig. 3a). During the second year the use of sally-strike maneuver is 90% lower than in the other two years, and the proportion of use of sally-hover maneuver is higher compared to the previous and subsequent years (Fig. 3a). In addition, an increase in the use of glean maneuver was observed towards the third year (Fig. 3a). The standardized Pearson residuals provide similar information: high absolute values for the sally-strike maneuver in the first year, and for the second-year high absolute values for sally-strike and sally-hover (Table 2). Greater Wagtail-tyrant and Ringed Warbling-Finch the frequency of use of the different maneuvers was independent of sampling year (Fig. 3b and c respectively).

**Table 2.** Pearson’s standardized residuals for the different categories of attack maneuvers and sampling years for Grey-crowned Tyrannulet inhabiting central Monte desert, Argentina. Cells with standardized residual values indicating a lack of fit to the hypothesis of independence in the cell are highlighted in bold.

| Sampling year | *Attack maneuvers* |  |  |
| --- | --- | --- | --- |
|  | Glean | Sally-hover | Sally-strike |
| Year 1 | -0.96 | -0.70 | **4.64** |
| Year 2 | -0.66 | **2.03** | **-2.69** |
| Year 3 | 1.93 | -1.14 | 0.12 |

**Figure 3.**
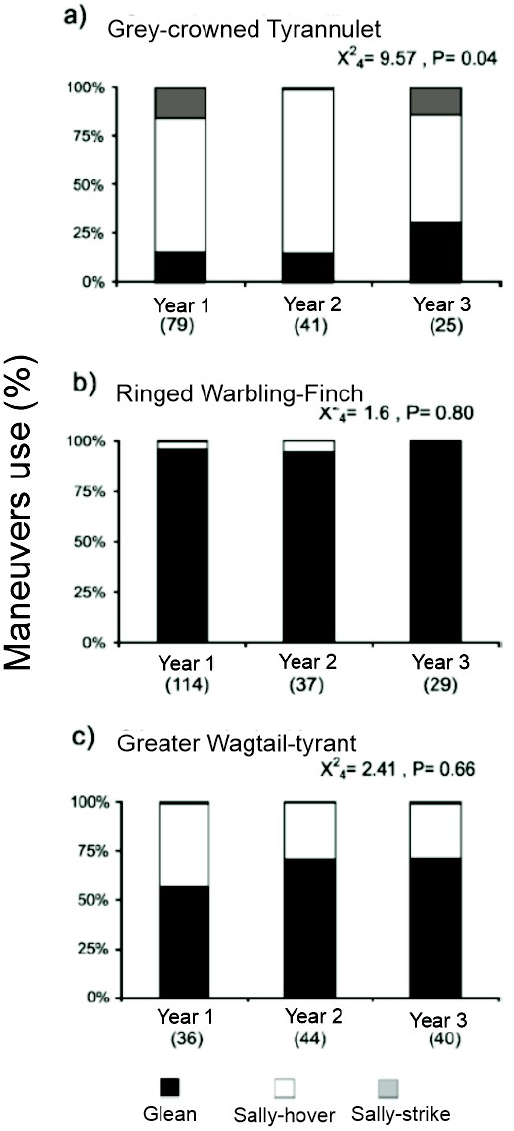
Prey-attack maneuvers used for three insectivorous bird species during three years in the central Monte desert, Argentina. The number of independent sequences with at least one attack (N) for each bird species each year is indicated in parentheses. The value of the Chi-square statistic and the P value for the test of independence are indicated.

### Arthropods abundance and biomass

Arthropod abundance per branch was statistically different among years (L= 46.02, gl=2, P<0.0001, log likelihood ratio test, comparing with the null model). All years were different from each other (year 1-year 2: z= −2.31, P=0.02; year 1-year 3: z= 3.7, P<0.001; year 2-year 3: z= 7.63, P<0.0001, Fig. 4) when looking at the statistics for each level of the “year” factor and comparing with the intercept of the generalized linear model using the Negative Binomial distribution. The lowest abundance of arthropods in branches corresponded to the second year and the highest abundance to the third year (Fig. 4).

**Figure 4.**
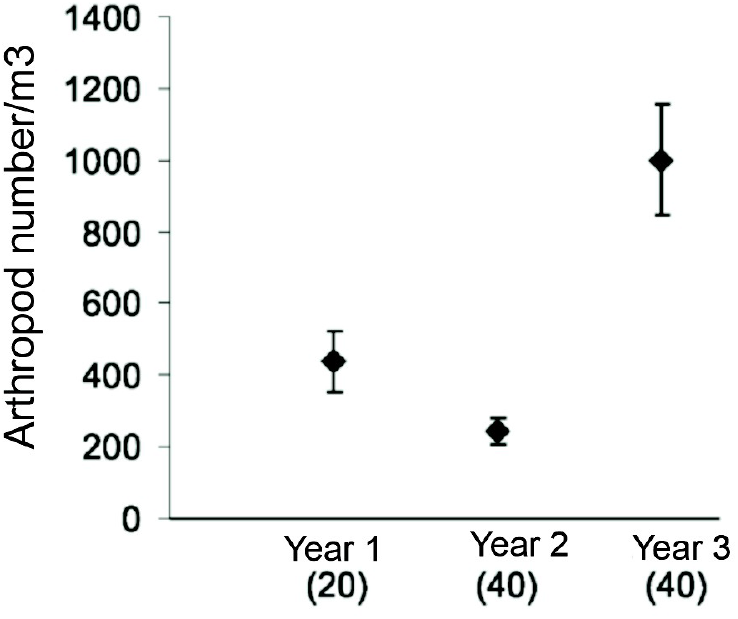
Arthropod abundance per cubic meter of *Neltuma flexuosa* branch during three years in the central Monte desert, Argentina. The number of samples for each year is indicated in parentheses. Although the number of arthropods per m^3^ is expressed graphically, the generalized linear model was performed with the variable Number of arthropods per branch with the volume of each branch as an offset variable (see Methods).

For the arthropod biomass analyses the heteroscedasticity produced by the categorical factor “year” was modeled. The comparison between the model without the incorporation of the variance structure and the model with the modeled heteroscedasticity proved to be significant (Table 3). Furthermore, from the comparison of the AIC statistics (Akaike’s information criterion) the best model included the modeled heteroscedasticity (Table 3). Biomass was statistically different for the three years (L=77.57, gl=4, P<0.0001, log likelihood ratio test, Fig. 5). Biomass was significantly lower for the second year compared to first (t=3.00, P<0.01) and third year of the study (t=3.9, P<0.001; Fig. 5), and it did not differ between the first and third year (t= −1.37, P=0.17; Fig. 5). Frequency distribution of arthropod biomass was dependent of the year (X^2^_10_ = 113.33, P <0.0001, Test of Independence). The differences among the three years are mainly related with a lower frequency of small arthropods during the first 1 and a higher number of individuals of these sizes in the third year (Fig. 6). Arthropods collected in the third year are highly concentrated in the first mass categories (mainly small dipteran larvae and 0.05 - 0.15 mg homopteran nymphs). In the following categories the pattern of frequencies is reversed for the three years, with first and second years having the highest percentage of heavy biomass arthropods. During these two years most of the individuals correspond to Homoptera of the superfamilies Jassoidea and Fulgoroidea and Coleoptera of the superfamily Curculionoidea with a biomass of approximately 0.5 - 1 mg; individuals of more than 10 mg (larvae of Leidoptera and Homoptera of the family Membracidae) were also recorded, while in the third 3 no individuals of more than 5 mg were recorded (Fig. 6).

**Table 3.** Comparison of the general linear model corresponding to the analysis of biomass per m^3^ of arthropods with its corresponding GLS where heteroscedasticity was modeled, incorporating a variance structure in the “year” factor. The models were compared using a log likelihood ratio test.

|  | *gl* | *AIC* | *Log likelihood* | *L. ratio (L)* | *P* |
| --- | --- | --- | --- | --- | --- |
| GLM <sup>a</sup> | 4 | 1334.7 | -663.3 |  |  |
| GLSM “year” <sup>b</sup> | 6 | 1280.7 | -634.3 | 57.9 | < 0.0001 |

<sup>a</sup> General linear model.
<sup>b</sup> Generalized least squares model with variance structure in the “year” factor.

**Figure 5.**
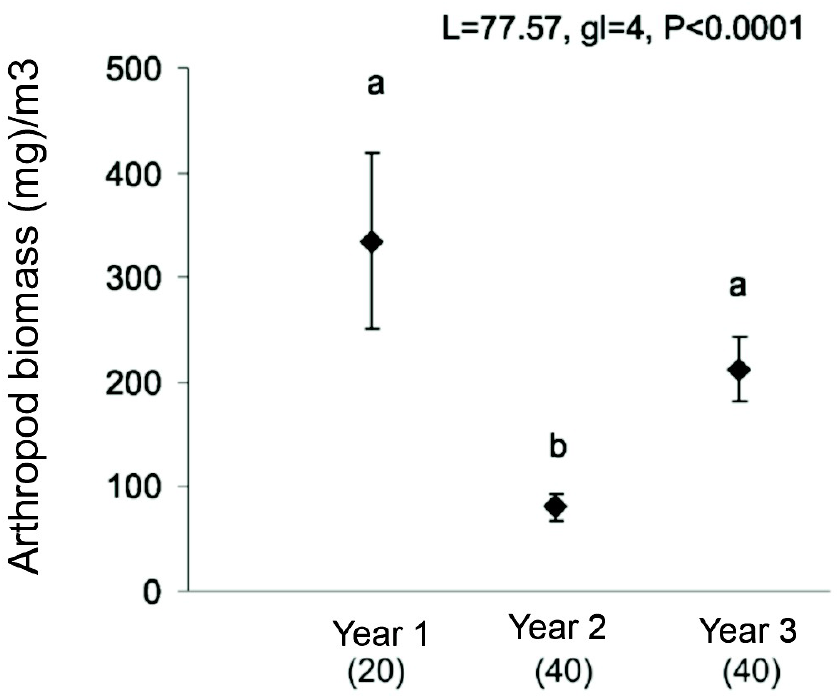
Arthropod biomass (mg) per cubic meter of *Neltuma flexuosa* branch (+ EE) for three years in the central Monte desert, Argentina. The number of samples for each year is indicated in parentheses.

**Figure 6.**
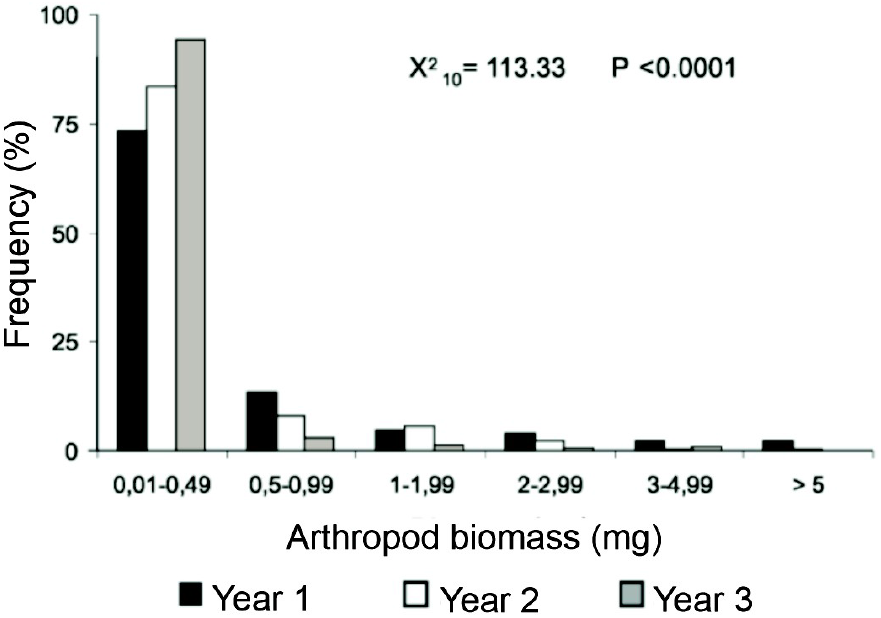
Frequency distribution of arthropod biomass (mg) collected on *Neltuma flexuosa* foliage during three years in the central Monte desert, Argentina. Pearson’s Chi-square statistic is indicated for the Test of Independence.

## Discussion

Factors that may explain interannual variation in bird foraging behavior include availability of food resources, climatic conditions, predation, and vegetation phenology (Szaro et al. 1990). During the study period of our research there were notable variations among years in the abundance and biomass of arthropods found in the foliage of *Neltuma flexuosa*, which could have affected the foraging behavior of birds, as has been observed in several works that studied this behavior interannually (Szaro et al. 1990, Bell and Ford 1990, Cueto et al. 2016).

Research of foraging behavior indicate that search speed may decrease in the following cases: a) when the abundance of more cryptic prey (Gendron 1982) and/or smaller prey (Goss-Custard 1977) increases, since fewer movements per unit of time improve the probability of prey detection, or b) when more prey is available (Zach and Falls 1979, Pienkowski 1983), since fewer search movements are necessary to find available prey. In our study, the three bird species maintained their search speed without variation, despite the fact that in the last sampling year the abundance of small prey increased. However, the three bird species changes their use of movements to search for prey and also Grey-crowned Tyrannulet change the maneuver used to capture preys. Searching and prey capture behavior of Grey-crowned Tyrannulet individuals corresponds to a variable distance searches foraging mode (Robinson and Holmes 1982), where birds perform more flight movements than jumps, and their main attack maneuver is sally-hover. However, Grey-crowned Tyrannulet changed their searching patterns from moving through flights in the first two years to moving using jumps to change perch as the main searching movement in the third year. This change in searching pattern was accompanied by an interannual change in the use of maneuvers to attack their prey, as in the third year decreased the use of sally-hover (their main maneuver in previous years) and increased the use of glean maneuver. These changes could be related to the type of prey more abundant in the foliage of *Neltuma flexuosa* during the third year of the study. In this year the abundance of arthropods was much higher than in previous ones, but most of them were of small size and biomass (mainly dipteran larvae and homopteran nymphs). For birds to capture this type of small prey the sally-hover maneuver might not be the most appropriate, probably because birds detect these preys better from a short distance; also moving with short movements (such as jumps) could allow birds to explore branches and foliage in greater detail. Therefore, being flexible in their feeding mode (and using another foraging mode such as “near surface searching”, Robinson and Holmes 1982) may be more appropriate during the third year. Similar changes to those occurring in Grey-crowned Tyrannulet have been observed in the foraging mode of Red-eyed Vireo (*Vireo olivaceus)*, this species generally possesses a “variable-distance searchers” foraging mode, however it used a foraging mode closer to the “near-surface search” syndrome in years in which there was an abundance explosion of lepidopteran larvae (Robinson and Holmes 1982). Other examples of foraging mode flexibility were found by Chen et al. 2011. These authors found that individuals of Yellow-billed Cuckoo (*Coccyzus americanus)* possess a combination of two feeding modes; individuals generally use the passive mode of foraging called “sit and wait” when there was large larvae (which can be detected at distances greater than 1 m and are captured by long jumps), however when these prey was not abundant, birds use the “wideling foraging” mode, which involves active foraging by short movements (Chen et al. 2011).

Interannual changes were also observed among the three years for Greater Wagtail-tyrant, and similar to the Grey-crowned Tyrannulet. Greater Wagtail-tyrant decrease in the use of flight searching movements and the perch changes by jumping during the second and third years in response to variations in the abundance and biomass of arthropods. In contrast, Ringed Warbling-Finch showed a very stereotyped behavior during the entire study period, using almost exclusively jumping and gleaning as a search and capture strategy. Further, during the third-year individuals of this species reinforcement their searching movements by jumping and decreased their search using flight movements. For Ringed Warbling-Finch, given the characteristics of its foraging behavior, the aforementioned changes in the abundance of the different types of arthropods did not particularly influence the way it attacks its prey, since it has the characteristics of a gleaner forager, which were in accordance with the changes in the abundance and size of the arthropods during the third year.

The interannual changes observed in searching and capture behavior provide valuable information on the degree of flexibility of birds in terms of their foraging modes, as well as the factors that could be influencing them. This ecological flexibility is of particular importance, as it could be advantageous for birds in environments with variable climates and food fluctuations (Morse 1980) such as desert environments.

